# Crystallography of lamin A facilitated by chimeric fusions

**DOI:** 10.1101/2020.02.28.969220

**Authors:** Giel Stalmans, Anastasia V. Lilina, Sergei V. Strelkov

## Abstract

All proteins of the intermediate filament (IF) family contain the signature central α-helical domain which forms a coiled-coil dimer. Because of its length, past structural studies relied on a ‘divide-and-conquer’ strategy whereby fragments of this domain were recombinantly produced, crystallized and analysed using X-rays. Here we describe a further development of this approach towards structural studies of nuclear IF protein lamin. To this end, we have fused lamin A fragments to short N- and C-terminal capping motifs which provide for the correct formation of parallel, in-register coiled-coil dimers. As the result, a chimeric construct containing lamin A residues 17-70 C-terminally capped by the Eb1 domain was solved to 1.83 Å resolution. Another chimera containing lamin A residues 327-403 N-terminally capped by the Gp7 domain was solved to 2.9 Å. In the latter case the capping motif was additionally modified to include a disulphide bridge at the dimer interface. We discuss multiple benefits of fusing coiled-coil dimers with such capping motifs, including a convenient crystallographic phasing by either molecular replacement or sulphur single-wavelength anomalous dispersion (S-SAD) measurements.

## Introduction

Lamins represent the fifth class of intermediate filament (IF) protein family. These nuclear proteins are expressed in all cell types and involved in a broad variety of cellular functions. A-type lamin (and its splice variant lamin C) as well as closely related lamin B1 and B2 jointly form the lamina, a meshwork of ∼3.5 nm thick filaments located at the inner side of the nuclear envelope (Turgay et al., 2017). Besides providing mechanical stability, lamins are involved in chromatin organization and transcription, DNA replication and repair, cell differentiation, mitosis and gene expression (Collas et al., 2014; Frock et al., 2006; Maynard et al., 2019; Qi et al., 2015; Shimi et al., 2008; Shumaker et al., 2008). Importantly, mutations in the lamin genes cause a wide variety of diseases, called laminopathies, and malfunctioning of lamins plays a role in diabetes, heat-shock and even cancer (Broers and Ramaekers, 2014; de Toledo et al., 2020; Pradhan et al., 2020).

Just like all IF members, the primary lamin structure consists of a central α-helical rod domain which is flanked by a non-helical N- and C-terminal domain (called head and tail, respectively). The central α-helical rod domain itself is divided into three segments known as coil 1A, coil 1B and coil2 which are interconnected through two linkers, L1 and L12 respectively. However, lamins own three unique regions in their primary structure: a 42-residue insert in coil1B (Lilina et al., 2020), a nuclear localization signal in the tail and a highly interacting s-type Ig-fold like motif in the tail. The basic unit of the lamin filament is a rod-like dimer, formed by two parallel monomers which create a coiled-coil (CC) structure covering the hole central α-helical rod domain. Moreover, this CC structure is characterized by an alternating pattern of hydrophobic and hydrophilic residues containing mostly 7-residue (heptad: a-g) repeats as well as several 11-residue (hendecad: a-k) repeats, in which positions a/d and a/d/h, respectively, are generally occupied by hydrophobic residues. Dependent on the repeat the CC structure is defined as left-handed (heptad) or essentially straight (hendecad) (Lupas and Bassler, 2017). Furthermore, the lamin CC structure has a 4-residue insert (stutter) located at the second part of coil2 (Strelkov and Burkhard, 2002).

Crystallographic studies have been essential towards the structural elucidation of lamin dimers. While the full-length dimer is too elongated and flexible to be crystallized, it was shown to be possible to crystallize and resolve fragmental structures (Strelkov et al., 2001). This ‘divide-and-conquer’ approach has helped to unravel great parts of the CC structure in vimentin, keratins as well as lamins to atomic resolution. Nevertheless, there still are missing gaps in the IF architectures, not only present at the dimer level but also present at higher assembly states.

One particular challenge linked to the use of shorter fragments of the IF rod domain is the correct formation of parallel, in register dimeric CCs. Such a structure may be reliably formed by the full-length protein (such as e.g. upon renaturation of recombinant IF proteins which are often purified from inclusion bodies (Herrmann et al., 2004)). However, the shorter fragments may not oligomerize at all, form CCs with wrong multiplicity such as trimers, or antiparallel and staggered structures rather than parallel and unstaggered. While also seen for numerous other CC proteins, this sort of complications were documented for multiple IF fragments designated for structural studies (Chernyatina et al., 2016). This was true in particular for the fragments corresponding to the N- and C-terminal parts of the rod domain. These parts are generally believed to be crucially involved in the longitudinal assembly of the filament (Herrmann et al., 2000; Strelkov et al., 2003). For instance, the isolated coil1A segment of vimentin was initially crystallized as a monomer (PDB code 1GK7), while it was later found out that it is only marginally stable as a dimer in solution (Meier et al., 2009; Strelkov et al., 2002). Another example is a short lamin A fragment (residues 328-398, PDB code 2XV5) that was found to engage in an unexpected, staggered assembly (Kapinos et al., 2011).

Here we have focused on two shorter fragments of lamin A (each of about 50 residues) which represent the N- and C-terminal portions of the rod domain. Such fragments (originally dubbed ‘minilamins’ (Kapinos et al., 2010)) were proposed towards atomic resolution studies of the interactions driving the longitudinal assembly of lamin dimers (Strelkov et al., 2004). It was shown that isolated N- and C-terminal fragments would indeed interact in solution (Kapinos et al., 2010). Specifically, we describe stabilization of both N- and C-terminal fragments by carefully fusing them to specific capping motifs which secure the formation of the correct CC dimer and are able to provide additional benefits for X-ray analysis. Such chimeras are successfully crystallized and resolved within this research as well as other research concerning tropomyosin and myosin (Frye et al., 2010; Korkmaz et al., 2016).

### Design of lamin A fragments for structural studies

Historically, stabilization of an IF protein fragment was first demonstrated using a 31 residue long GCN4 leucine zipper which is known as a prototype CC dimer featuring high thermal stability (O’Shea et al., 1991). To this end, a chimeric construct containing the GCN4 zipper followed by the last 28 residues (385-412) of the rod domain of human vimentin was created (Strelkov et al., 2002). Despite this particular design being successful in the sense of enabling crystallographic studies, fusing together two plain heptad-based sequences appears potentially troublesome, since wrong multiplicity or register of the CC are still possible when not designed carefully (Andreas et al., 2017). The most suitable solution appears to be a specific capping motif (N- or C-terminal) which would bring together the corresponding ends of the sequence of interest, ‘bootstrapping’ the formation of the CC. A natural example here is the C-terminal motif, known as the ‘foldon’, of the trimeric CC protein fibritin from bacteriophage T4 (Tao et al., 1997). For stabilizing the dimeric CC fragments of tropomyosin and muscle myosin, bacteriophage ϕ29 scaffolding protein Gp7 (Morais et al., 2003) and microtubule binding protein Eb1 (Slep et al., 2005) domains were used in the past as N- and C-terminal caps respectively. These capping motifs have enabled X-ray structural determination of several myosin fragments as well as tropomyosin overlap (Frye et al., 2010; Taylor et al., 2015; Korkmaz et al., 2016). Both motifs are organized in a similar way. They are small (51 and 36 residues respectively), dimeric and feature, per monomer, a short α-helix engaging in the CC interface and an additional α-helix folding back onto the CC. Such motifs are very well suited to support a CC in the fragment of interest, provided the fusion preserves the hydrophobic heptad pattern.

The first construct created corresponds to residues 17-70 of human lamin A fused to residues 215-251 of Eb1. This construct includes the predicted coil1A domain of lamin (residues 28-67). The fusion is performed in such a way that a regular heptad repeat pattern is preserved from lamin coil1A into the Eb1 cap (Fig. 1A). The second construct involves residues 1-49 of Gp7 followed by residues 327-403 of human lamin A. It thus includes the last portion of coil2 which is predicted to terminate near residue 380. Importantly, the residues 327-330 (L-A-R-E) correspond to a stutter insert and has been preserved when creating the fusion (Fig. 1B).

**Figure 1.**
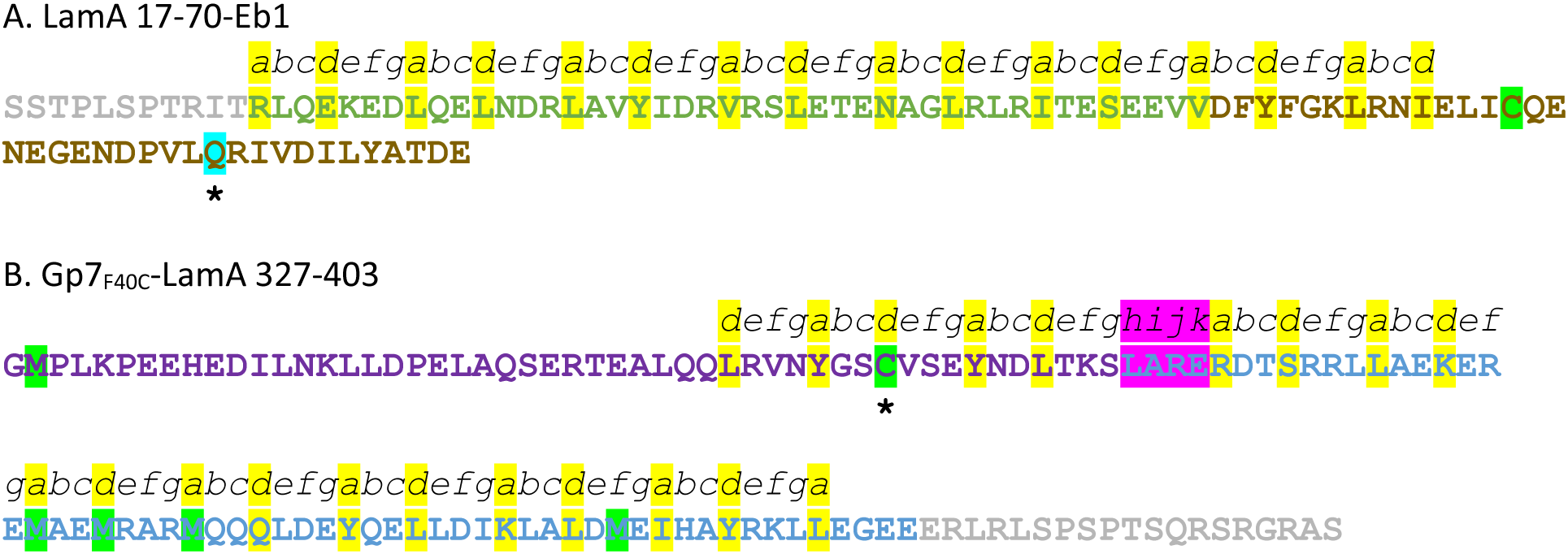
Amino acid sequences of the two fusion constructs. A. LamA 17-70-Eb1 construct. The heptad repeat pattern is highlighted yellow. The residue Gln that was mutated to Trp to enable crystal detection using fluorescence imaging is highlighted in cyan. The natively present Cys residue is highlighted in green. The lamin A region is coloured grey (head) and green (coil 1A) whereas the Eb1 cap is coloured brown. B. Gp7_F40C_-LamA-327-403. The heptad repeat pattern is highlighted yellow. The Cys residue introduced by point mutation to enable disulphide formation is highlighted in cyan. The natively present Met residues are highlighted in green as well. The stutter region is highlighted in purple. The lamin A region is coloured blue (coil 2) and grey (tail) whereas the Gp7F40C cap is coloured deep purple.

Another key benefit of including a capping motif towards X-ray analysis is the possibility to use this motif as (part of) the starting model during the initial phasing of crystallographic data through molecular replacement (MR) (Evans and McCoy, 2007). While MR is often possible for a simple CC, the success of this procedure varies greatly depending on multiple factors including the length of the CC, space group symmetry, packing density etc. (Guzenko et al., 2017; Thomas et al., 2015). These difficulties are largely attributed to the very nature of CCs: in particular, a typical heptad-based structure overlaps on itself upon a seven-residue register shift. By including a capping motif at one end of a CC it is possible to reduce this internal symmetry and thus increase the chances towards a successful MR search.

In addition, a further, independent aid in the crystallographic phasing was provided through a point mutation of a Phe residue (F40) within the CC stalk of the Gp7 cap to a Cys residue (Fig. 1B). This particular cysteine was introduced in ‘d’ position of the CC. Previously we have demonstrated that this heptad position is well suited for the formation of a disulphide (Chernyatina and Strelkov, 2012). Indeed, this could be eventually confirmed for the Gp7_F40C_-LamA 327-403 chimera using non-reducing SDS-PAGE (data not shown). For experimental phasing using single-wavelength anomalous dispersion on sulphur atoms, a disulphide is highly beneficial as it produces a more reliable signal compared to single sulphurs. The utility of disulphides for the phasing of CC structures in particular has recently been demonstrated (Kraatz et al., 2018).

Likewise, independent aid in the crystallization screening was provided through a point mutation of a Gln residue (Q26) within the back-folding CC of the Eb1 cap to a Trp residue. This point mutation enabled fluorescence imaging (UVEX-PSA42™ (Jan Scientific, USA)) of LamA 1-292-Eb1_Q26W_ crystals (non-described chimera) that initially had no tryptophan or tyrosine (data not shown). Nevertheless, due to chemical environment and composition of the crystallization buffer native fluorescence imaging can fail. Luckily this can be overcome by using 2,2,2-trichloroethanol as fluorescence enhancer, thereby not impairing the diffraction quality of crystals and having no significant effect on crystal structure as shown by Pichlo et al (Pichlo et al., 2018).

### Cloning, expression, purification and crystallization screening

The overall strategy towards the production of chimeras was as described by Chernyatina et al (Chernyatina et al., 2016). Initially the DNA sequence for the capping motifs were purchased as gBlocks Gene Fragments from Integrated DNA Technologies. Prior to cloning, a Quick-Change™ site-directed mutagenesis was performed to introduce a cysteine (F40C) into the Gp7 cap. A Sequence Ligation Independent Cloning was performed using a pETSUK2 vector (Weeks et al., 2007). The resulting plasmids encoded a 6xHis tag and a small ubiquitin related modifier (SUMO) cleavage site at the Lamin A N-terminus.

Overexpression was done in an E. Coli Rosetta 2 (DE3) pLysS strain (Merck, Germany) by auto-induction based on the ZYP-5052 medium (Studier, 2014, 2005). The cells were harvested by centrifugation, resuspended in low-imidazole buffer (12.5 mM imidazole, 250 mM NaCl, 5 mM βME, 40 mM Tris pH 7.5) containing a lysis mixture, sonicated and clarified by centrifugation. The supernatant was loaded onto a Ni-column (His60 Ni Superflow Resin (Takara Bio, USA)), pre-equilibrated with low-imidazole buffer. 6xHis-SUMO tagged chimeric fusions were eluted by applying high-imidazole buffer (500 mM imidazole, 250 mM NaCl, 5 mM βME, 40 mM Tris pH 7.5). The 6xHis-SUMO tag was cleaved by overnight incubation with SUMO Hydrolase 7K while dialyzing against low-imidazole buffer. Afterwards, the sample was re-loaded onto the Ni-column, pre-equilibrated with low-imidazole buffer. The chimeric fusions were eluted with low-imidazole buffer while 6xHis-SUMO tags were trapped on the Ni-column. The final purification step was gel-filtration on a Superdex 200 Increase column (GE Healthcare Europe, Belgium) in 10 mM Tris pH 7.5, 150 mM NaCl.

The purified fractions containing chimeric fusions were concentrated (8.14 mg/mL, LamA 17-70-Eb1 and 9.3 mg/mL, Gp7_F40C_-LamA 327-403) using Amicon^®^ Ultra filters with 3kDa cut-off (Merck Millipore, Belgium) extensively screened for crystallization and optimized until a satisfied diffraction level was reached.

### Crystallographic structure determination

Purified LamA 17-70-Eb1 (8.14 mg/mL) was crystallized at 4°C by sitting drop vapour diffusion method using 35.7% (w/v) 1,6-hexanediol, 5% (w/v) PEG 1K and 0.1 M trisodium citrate dihydrate (pH 4.9). Crystals were mounted on cryo-loops using mother liquor supplemented with 30% (v/v) glycerol. Native data were collected at beamline I04, Diamond Light Source (UK). Standard processing in XDS (Kabsch, 2010) was performed to obtain diffraction data up to 1.83 Å resolution. An initial model was obtained using molecular replacement by means of the Eb1 cap (dimeric form) as search model in Molrep (Vagin and Teplyakov, 2010).

Purified Gp7_F40C_-LamA 327-403 (9.13 mg/mL) was crystallized at 4°C by hanging drop vapour diffusion method using 35% (v/v) methanol, 0.2 M MgCl_2_ and 0.1 M HEPES (pH 8.2). Crystals were mounted on cryo-loops using mother liquor supplemented with 30% (v/v) glycerol. Native data were collected at beamline Proxima-1, Synchrotron Soleil (France). Standard processing in XDS was performed to obtain a complete diffraction data set up to 2.9 Å resolution. Additional data were collected using the wavelength 1.8 Å with a redundancy of 120 towards phasing by anomalous dispersion. The high-redundancy long-wavelength dataset has revealed a significant anomalous signal up to 3.2 Å resolution. The data were processed through the Auto-Rickshaw web-server (Panjikar et al., 2005). Initially, SHELXD (Sheldrick, 2008) showed 21 heavy atom sites above 0.2 occupancy per monomer. Further processing by Phaser (Read and McCoy, 2011) eventually determined the position of 24 anomalous peaks per dimer. This included all 12 sulphur atoms in the dimer (the disulphide bridge; methionine cluster consisting of Met345, Met349 and Met352; Met278 and Met371). The first traced position was the disulphide bridge formed by F40C mutation in the Gp7 cap, the second one was two Met351 residues in the centre of the dimer (Fig. 4). Interestingly, Auto-Rickshaw also indicated the presence of heavy atom peaks that were not sulphur but rather heavy-atoms bound during purification procedure (i.e. position #4 that can be Ni coordinated by two His285 (Fig. 4)). Determining the position of the heavy atoms enabled experimental phasing of the complete structure. The model was further improved using Buccaneer (Cowtan, 2006) that was also part of the Auto-Rickshaw pipeline. The initial maps showed the placement of the Gp_740C_ structure that helped to determine the right orientation of the lamin CC.

**Figure 2.**
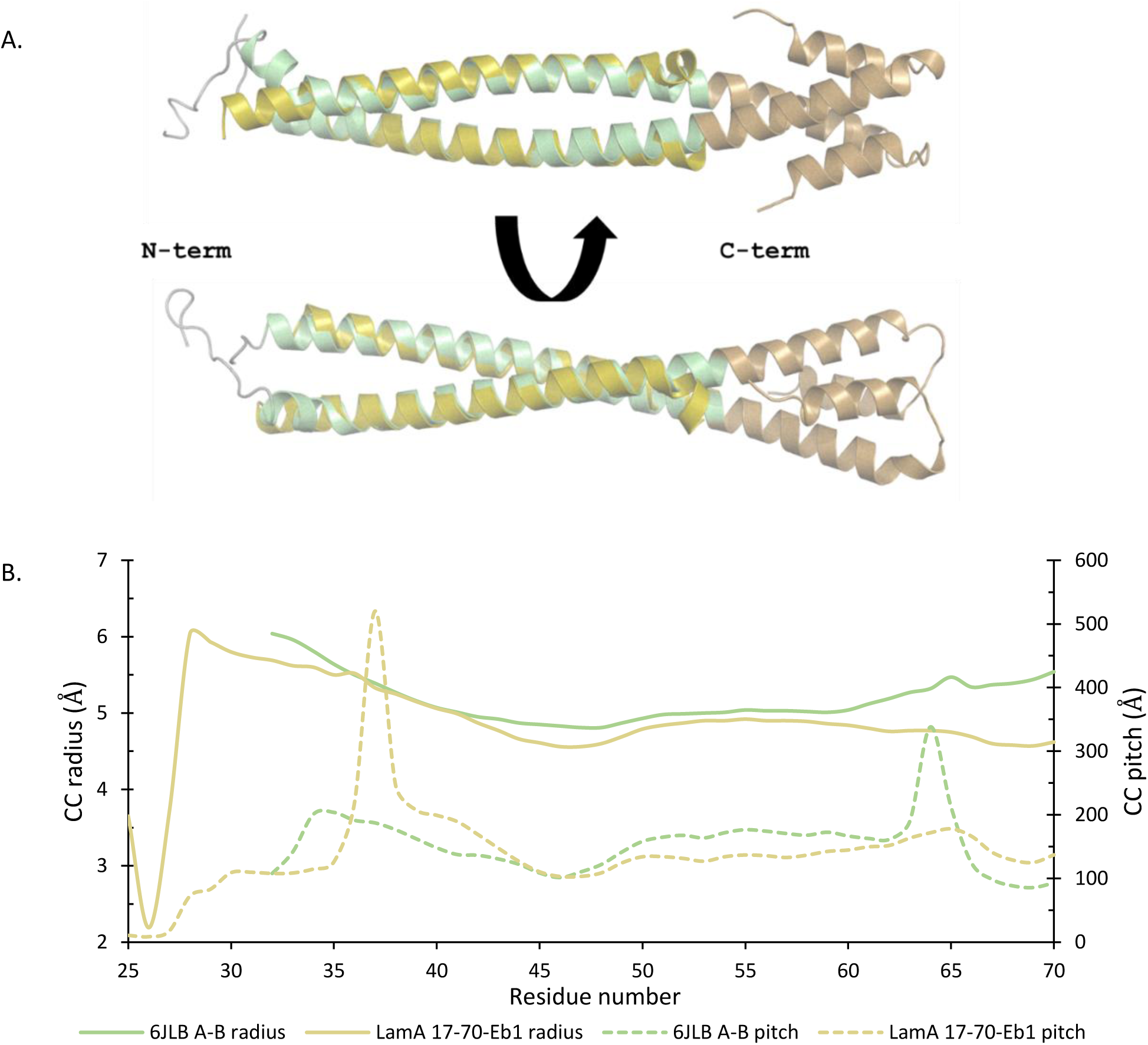
A. Coil1A dimer superposition of the 6JLB A-B (gold) and LamA 17-70-Eb1 (grey-green-brown) atomic structures. B. Corresponding plots of the coiled-coil radius and pitch.

**Figure 3.**
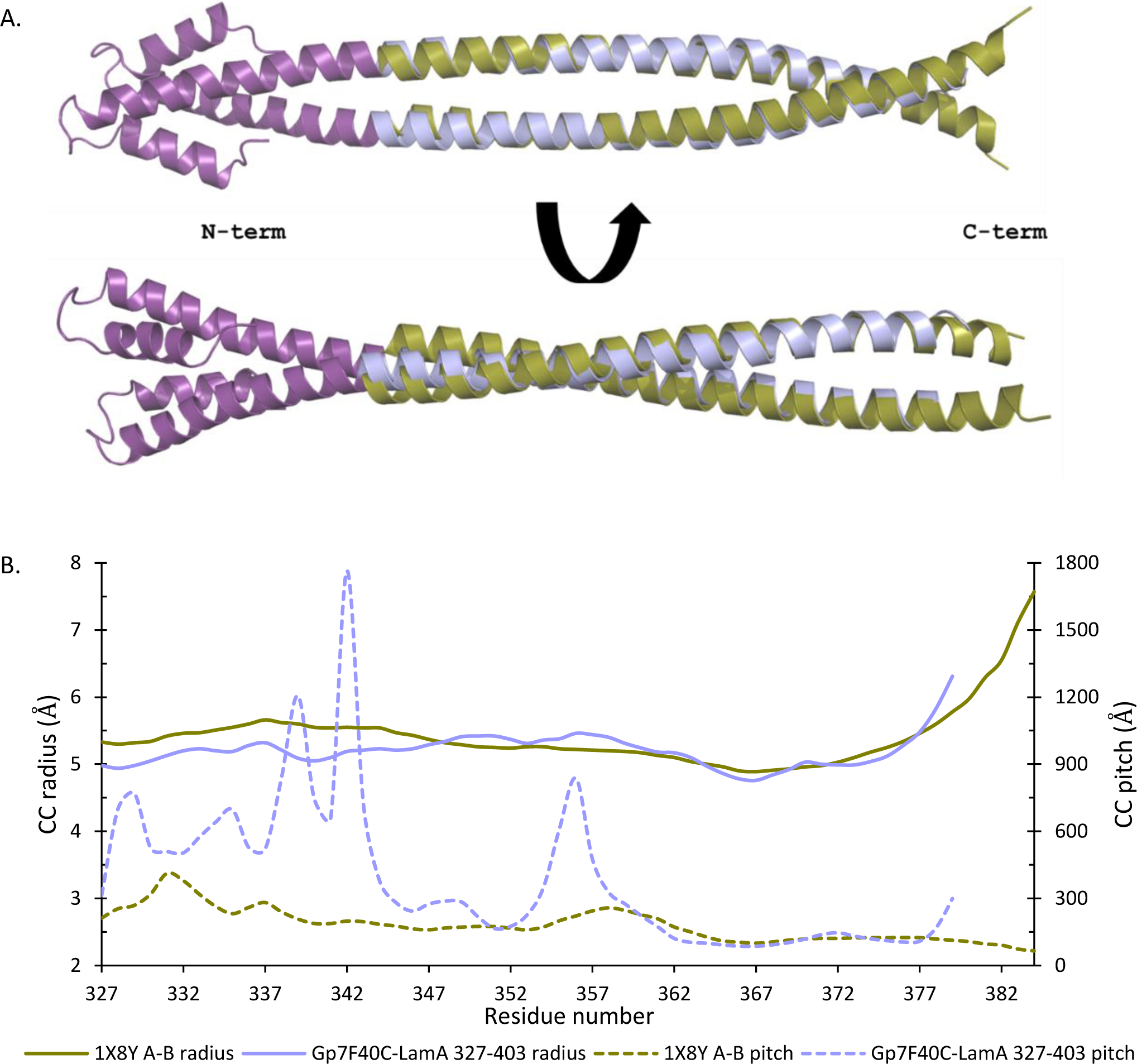
A. Coil2 dimer superposition of the 1X8Y A-B (olive) and Gp7_F40C_-LamA 327-403 (purple-blue) atomic structures. B. Corresponding plots of the coiled-coil radius and pitch.

**Figure 4.**
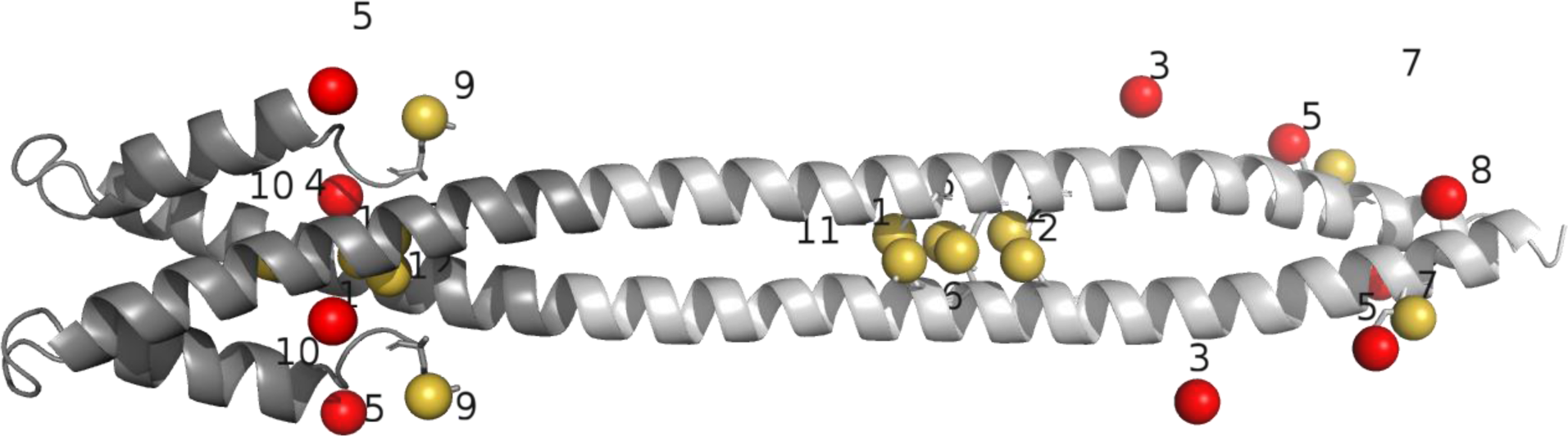
Atomic structure of the Gp7_F40C_-LamA 327-403 construct (grey) with heavy atom sites. Sulphur atoms present in the structure are shown as yellow spheres, while red spheres represent other suggested sites. Numbers show the order in which the sites were identified by Phaser (Read and McCoy, 2011), as part of the Auto-Rickshaw pipeline (Panjikar et al., 2005).

For both structures, Coot was used for initial real-space refinement and automated refinement was carried out using Refmac5 (Emsley et al., 2010; Vagin et al., 2004). Regarding the Gp7_F40C_-LamA 327-403 structure, Buster was also used for automated refinement (Bricogne et al., 2017). CC geometry was analysed using the TWISTER program (https://pharm.kuleuven.be/apps/biocryst/twister.php, Strelkov and Burkhard, 2002). Furthermore, Pymol (The PyMOL Molecular Graphics System, Version 2.0 Schrödinger, LLC) was used to prepare figures for both chimeric fusions. All details on crystallographic structure determination are shown in Table 1.

**Table 1.** Information on crystallographic structure determination of LamA 17-70-Eb1 and Gp7_F40C_-LamA 327-403. Sequence data in square brackets are for the capping motifs alone. Crystallographic data in round brackets are for the highest resolution shell.

|  | **LAMA 17-70-EB1**<br>**[EB1 CAPPING MOTIF ONLY]** | **GP7<sub>F40C</sub>-LAMA 327-403**<br>**[GP7<sub>F40C</sub> CAPPING MOTIF ONLY]** |  |
| --- | --- | --- | --- |
| **Sequence features** |  |  |  |
| No. of residues | 91 [35] | 127[49] |  |
| No. of Cys residues | 1 [1] | 1 [1] |  |
| No. of disulphides per dimer | 0 [0] | 1 [1] |  |
| No. of Met residues | 0 [0] | 5 [1] |  |
| **Diffraction data** |  | **Native** | **Anomalous** |
| Number of crystals used | 1 | 1 |  |
| Number of datasets collected | 1 | 1 | 3 |
| Space group | P 2<sub>1</sub> 2<sub>1</sub> 2<sub>1</sub> | P 6<sub>1</sub> 2 2 |  |
| Unit cell dimensions: |  |  |  |
| a, b, c (Å) | 27.37, 105.24, 119.77 | 117.25, 117.25, 93.156 |  |
| α, β, γ (°) | 90, 90, 90 | 90, 90, 120 |  |
| Resolution range (Å) | 48.2-1.8 (1.9-1.8) | 44.6-2.9 (3.0-2.9) | 49.7-3.2 (3.3-3.2) |
| <I/σ> | 9.27 (0.60) | 20.27 (0.99) | 60.19 (5.95) |
| CC<sub>½</sub> (%) | 0.999 (-0.311) | 0.999 (0.934) | 1 (0.998) |
| Completeness (%) | 99.92 (99.90) | 98.28 (98.24) | 98.06 (94.26) |
| No. of unique reflections | 27312 (3052) | 8792 (847) | 11862 (1181) |
| Redundancy | 10.4 (8.0) | 19.2 (20.1) | 122.2 (84.3) |
| Wilson B-factor (Å<sup>2</sup>) | 26.18 | 110.58 | 129.51 |
| **Refinement** |  |  |  |
| R<sub>work</sub> (%) | 22.4 | 26.7 |  |
| R<sub>free</sub> (%) | 25.6 | 32.6 |  |
| No. of protein chains per a.s.u. | 4 | 1 |  |
| No. of non-hydrogen atoms: |  |  |  |
| protein | 2638 | 850 |  |
| ligands/ions | 0 | 11 |  |
| water molecules | 97 | 1 |  |
| R.m.s. deviations: |  |  |  |
| bonds (Å) | 0.017 | 0.015 |  |
| angles (°) | 2.04 | 2.23 |  |
| Ramachandran plot:<br>favored/allowed/outlier<br>residues (%) | 99.4/0.6/0 | 96.1/2.9/1 |  |
| Side-chain rotamer outliers (%) | 2.4 | 12 |  |
| Clash score | 6.33 | 5.24 |  |

### Structural features and conclusions

The first fusion construct LamA 17-70-Eb1 could be readily superposed to the coil1A dimer resolved within the recently published X-ray structure of lamin A fragment encompassing residues 1-300 at 3.2 Å resolution (PDB code 6JLB A-B, Ahn et al., 2019). The superposition (Fig. 2A) reveals an identical regular left-handed CC structure (Fig. 2B), with a root mean square deviation of 0.935 Å for 84 Cα-positions. While in the lamin A 1-300 structure the first ∼30 residues corresponding to the head domain are not resolved, the superior diffraction quality of the LamA 17-70-Eb1 crystals has helped to trace the structure starting with residue 24 in both chains.

Likewise, the second fusion construct Gp7_F40C_-LamA 327-403 superposes well (Fig. 3A) with the previously resolved structure of lamin A fragment 305-387 (PDB code 1X8Y, Strelkov et al., 2004), with a root mean square deviation of 1.294 Å for 108 Cα-positions. It should be noted that the latter structure, despite a higher resolution (2.2 Å), suffered from an increasing disorder towards the N-terminal end. Our fusion with the Gp_740C_ cap provides for a proper stabilization of the N-terminal part of the lamin A sequence. As a result, the geometry of the CC formed by the chimera reveals a distinct unwinding (i.e. loss of the left-handed twist) near the stutter insert (Fig. 3B), in line with the theoretical expectations (Guzenko and Strelkov, 2018; Strelkov and Burkhard, 2002).

In conclusion, we have demonstrated that fusions of ∼50 residue long CC fragments from human lamin A with well-chosen N- and C-terminal capping motifs provides a number of benefits. First, it provides for the correct assembly of the dimeric, registered CC. Second, the addition of the capping motifs provides additional options for crystallographic phasing either by MR or by using the anomalous signal of sulphurs. It should also be noted that the capping motifs used are involved in the crystal contacts in both cases. We are currently exploring the benefit of using these capping motifs towards crystallization of longer CCs. Additional modifications within the capping motifs that would further enhance their utility are certainly possible. For instance, one could introduce further point mutations enabling additional disulphides on the CC axis or facilitating crystallization screening. In addition, it is feasible to couple additional groups at the free ends of the capping motifs that would render further functionalities. For instance, a motif for a heavy atom attachment would enable a straightforward phasing by anomalous signal, etc.

## Abbreviations

IF: intermediate filament;
CC: coiled-coil;
βME: β-mercaptoethanol;
SUMO: small ubiquitin related modifier;
MR: molecular replacement

## Acknowledgements

We thank the teams of the beamline I04 at Diamond Light Source (UK) and of the beamline Proxima-1 at Synchrotron Soleil (France) as well as Dr. Evgenii Osipov for help with crystallographic data collection.

